# Obesity-linked PPARγ S273 phosphorylation promotes insulin resistance through Growth Differentiation Factor 3

**DOI:** 10.1101/2020.01.13.904953

**Authors:** Jessica A. Hall, Deepti Ramachandran, Hyun C. Roh, Joanna R. DiSpirito, Thiago Belchior, Peter-James H. Zushin, Colin J. Palmer, Shangyu Hong, Amir I. Mina, Bingyang Liu, Zhaoming Deng, Pratik Aryal, Christopher Jacobs, Danielle Tenen, Chester W. Brown, Julia F. Charles, Gerald I. Shulman, Barbara B. Kahn, Linus T.Y. Tsai, Evan D. Rosen, Bruce M. Spiegelman, Alexander S. Banks

**Affiliations:** Division of Endocrinology, Diabetes and Metabolism, Beth Israel Deaconess Medical Center and Harvard Medical School, Boston, MA; Department of Immunology, Harvard Medical School, Boston, MA; Department of Pediatrics, University of Tennessee Health Science Center, Memphis, Memphis, TN; Department of Orthopedics, Brigham and Women’s Hospital, and Harvard Medical School Boston, MA; Department of Internal Medicine, Yale University School of Medicine, New Haven, CT; Dana-Farber Cancer Institute, Department of Cell Biology, Harvard Medical School, Boston, MA

**Author notes:** equal contributions (Jessica A. Hall, Deepti Ramachandran).

## Abstract

Overnutrition and obesity promote adipose tissue dysfunction, often leading to systemic insulin resistance. The thiazolidinediones (TZDs) are a potent class of insulin-sensitizing drugs and ligands of PPARγ that improve insulin sensitivity, but their use is limited due to significant side effects. Recently, we demonstrated a mechanism by which TZDs improve insulin sensitivity distinct from receptor agonism and adipogenesis: reversal of obesity-linked phosphorylation of PPARγ at Serine 273. However, the role of this modification has not been tested genetically. Here we demonstrate that mice encoding an allele of PPARγ which cannot be phosphorylated at S273 are protected from insulin resistance, without exhibiting differences in body weight or TZD-associated side effects. Indeed, hyperinsulinemic-euglycemic clamp experiments confirm improved insulin sensitivity, as evidenced by increased whole-body glucose uptake. RNA-seq experiments reveal PPARγ S273 phosphorylation specifically enhances transcription of Gdf3, a BMP family member. Ectopic expression of Gdf3 is sufficient to induce insulin resistance in lean, healthy mice. We find that Gdf3 can impact metabolism by inhibition of BMP signaling. Together, these results highlight the diabetogenic role of PPARγ S273 phosphorylation and focuses attention on a putative target, Gdf3.

## Introduction

Type 2 diabetes (T2D) is one of obesity’s most tightly linked comorbidities, and its prevalence is rising in parallel with the global obesity epidemic [1]. T2D results from elevated glycemia consequent to insulin resistance and beta cell failure [2]. Although there are several available glucose-lowering agents, few address the underlying pathophysiology of this disease. The thiazolidinediones (TZDs) are a class of insulin-sensitizing anti-diabetic drugs capable of delaying or even preventing the onset of diabetes [3]. Despite potent insulin sensitization, the clinical use of TZDs has declined due to adverse side effects [4]. As ligands for the nuclear receptor peroxisome proliferator activated receptor-γ (PPARγ), the therapeutic actions of TZDs have been assumed to result from their classical agonism towards PPARγ. This agonism drives adipocyte differentiation and lipogenesis in fat tissues [5]. PPARγ is most highly expressed in adipose tissue, where it was first characterized as a master regulator of adipocyte differentiation and function [6]. Nonetheless, activation of PPARγ improves insulin sensitivity owing to its diverse roles in both adipocyte and non-adipocyte cells, including macrophages, T-cells, myocytes, and others [7–10]. PPARγ is intimately linked to insulin sensitivity, and genetic variants in the *PPARG* locus associate with diabetes risk [11–13].

We have recently shown that, in addition to PPARγ agonism, TZDs have a second biochemical role, blocking phosphorylation on PPARγ at serine 273 [14]. Importantly, phosphorylation of this residue in adipose tissue increases in murine models of obesity, and its reversal correlates with the anti-diabetic effects of TZD administration in obese patients and rodents [14, 15]. Moreover, synthetic ligands have been generated that prevent PPARγ S273 phosphorylation in the absence of PPARγ agonism. These non-agonist PPARγ ligands exhibit profound glucose-lowering effects with reduced side effects, such as weight gain and fluid retention [16–18]. Likewise, inhibition of the major protein kinase responsible for PPARγ S273 phosphorylation, extracellular signal-regulated kinase (ERK), improves insulin sensitivity in obese animals [19]. Understanding the specific mechanisms linking insulin sensitivity with reversal of obesity-linked PPARγ S273 phosphorylation is thus attractive as a new therapeutic strategy for type 2 diabetes, as well as other conditions linked to insulin-resistance such as Polycystic Ovarian Syndrome.

Here, we specifically address the contribution of PPARγ S273 phosphorylation to the development of insulin resistance, using a genetically modified mouse model where PPARγ cannot be phosphorylated at S273. These data demonstrate that genetic prevention of PPARγ S273 phosphorylation is an insulin-sensitizing event and is associated with decreased levels of Gdf3 in adipose tissues and skeletal muscle. Gdf3 is therefore a biomarker of PPARγ S273 phosphorylation; furthermore, neutralization of Gdf3 is a potential insulin-sensitizing therapeutic approach.

## Results

### Improved insulin sensitivity in mice with non-phosphorylatable PPARγS273

Reversal of obesity-linked PPARγ S273 phosphorylation with small-molecule PPARγ ligands correlates with anti-diabetic properties in humans and mice. However, it is unknown whether genetic modification of this site will result in chronic improvements in insulin sensitivity. To test this, we have generated a genetic mouse model in which PPARγ cannot be phosphorylated at S273. Using homologous recombination, a single nucleotide was changed to cause a serine to alanine substitution at codon 273 of PPARγ (Suppl. Fig. 1A, 1B). Homozygous knock-in mice (PPARγ^S273A/S273A^, hereafter referred to as PPARγ^A/A^) were generated using C57BL/6 ES cells and were born at the expected Mendelian frequencies. Whereas wild-type (WT) mice made obese by exposure to a high fat diet contain PPARγ that is phosphorylated at S273 in adipose tissue, PPARγ from PPARγ^A/A^ mice is exclusively unphosphorylated (Suppl. Fig. 1C). PPARγ S273 phosphorylation increases with progressive obesity [14]. Consequently, WT mice are predicted to have little PPARγ phosphorylation in the lean state, rendering them phenotypically identical to lean PPARγ^A/A^ mice. On a standard chow diet, PPARγ^A/A^ mice gain weight similarly, have comparable body compositions, and respond similarly to WT controls in glucose tolerance and insulin tolerance tests (Suppl. Fig. 2A-D). Accordingly, in the absence of stimuli to promote PPARγ phosphorylation, differentiation of primary adipocytes is unchanged between genotypes (Suppl. Fig. 3A). Gene expression of WT and PPARγ^A/A^ primary adipocytes revealed similar mRNA levels of the differentiation markers *Pparg2, Fabp4, Cebpa, Lpl,* and *Cd36* (Suppl. Fig. 3B). These data confirm earlier work that modulation of PPARγ S273 phosphorylation is neither necessary nor sufficient to affect adipogenesis. We also examined baseline differences in expression of a gene set previously identified using PPARγ^-/-^ cells reconstituted with viruses expressing WT or S273A PPARγ. Surprisingly, expression of these genes was not significantly different in primary adipocytes differentiated *ex vivo* from these mice (Suppl. Fig. 3C).

Given that PPARγ S273 is phosphorylated in the setting of high-fat feeding [14], we maintained PPARγ^A/A^ and WT littermate mice on a high-fat diet (HFD) to promote obesity and insulin resistance. PPARγ^A/A^ and WT mice experienced equivalent weight gain (Fig. 1A) with similar fat and lean body mass composition (Fig. 1B). As PPARγ activation with TZDs is associated with weight gain [20, 21], we assessed energy balance by indirect calorimetry experiments in PPARγ^A/A^ mice. There were no significant differences observed for VO_2_, VCO_2_, energy expenditure, food intake, body temperature or overall energy balance between genotypes. We did observe a small but significant decrease in RER in the PPARγ^A/A^ animals, which is suggestive of increased fatty acid oxidation (Suppl. Figs 4A and 4B).

**Figure 1.**
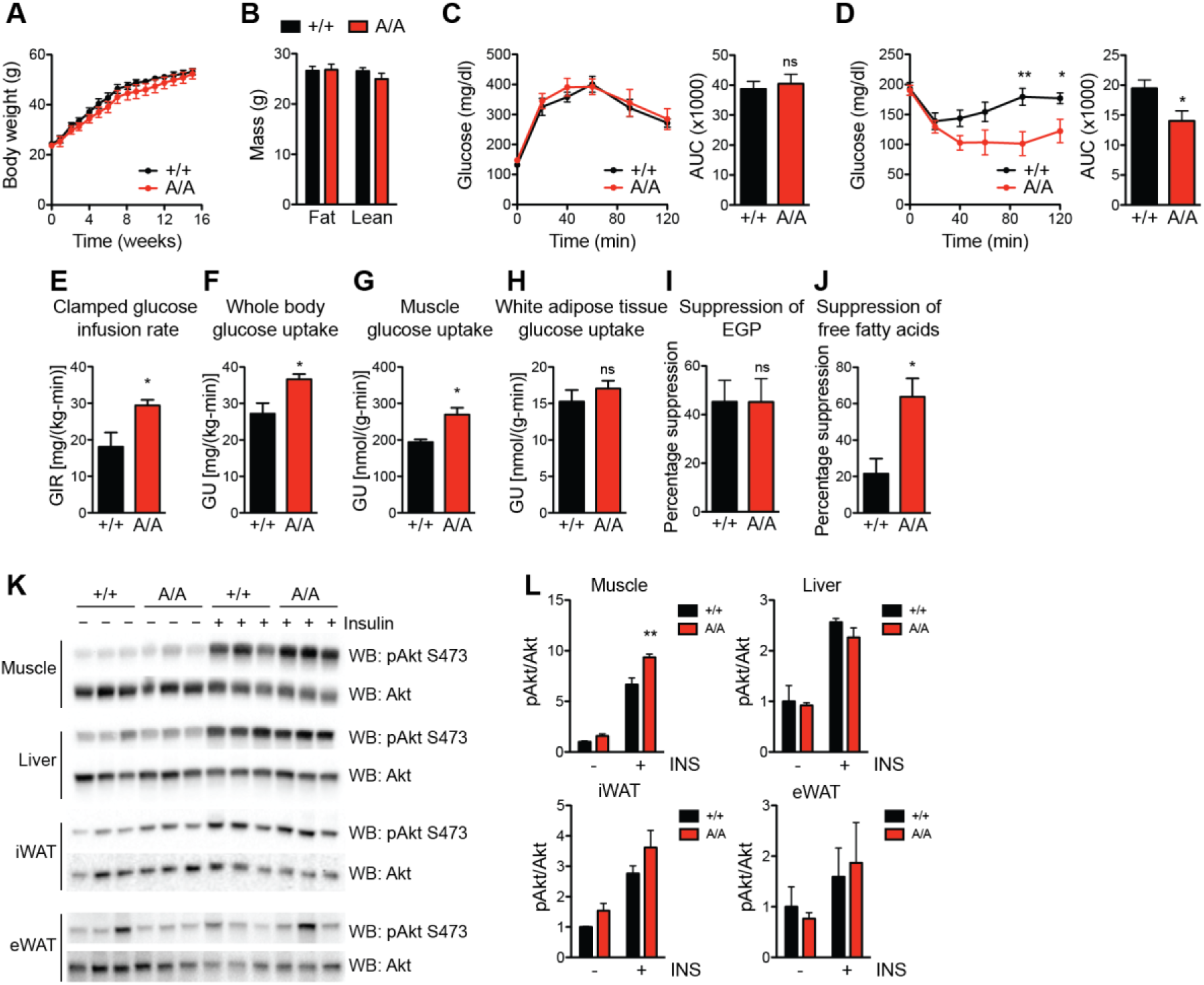
Improved insulin sensitivity in mice with non-phosphorylatable PPARγS273. Male wildtype (+/+) and PPARγA/A (A/A) mice were fed high-fat diet (HFD) for 16 weeks. A. Body weight of mice during HFD feeding (n = 6 +/+, 5 A/A). B. Body composition of mice at 16 weeks of HFD feeding. C. Glucose tolerance test (1 g glucose/kg body weight) after 15 weeks on HFD. D. Insulin tolerance test (2 U insulin/kg body weight) after 16 weeks on HFD. C-D. Right, area under the curve (AUC) for the respective test. Results are representative of three independent experiments. E-J. Hyperinsulinemic-euglycemic clamps in HFD-fed mice (n = 4 +/+, 5 A/A). E. Glucose infusion rate (GIR). F. Whole body glucose uptake (GU). G. Glucose uptake into skeletal muscle. H. Glucose uptake into epididymal white adipose tissue. I. Percentage suppression of endogenous glucose production (EGP). J. Percentage suppression of free fatty acids. K-L. Insulin sensitivity as determined by insulin-stimulated Akt phosphorylation (Ser 473) in tissues of HFD-fed wildtype and A/A mice. K. Immunoblot analyses of skeletal muscle, liver, inguinal white adipose tissue (iWAT), and epididymal white adipose tissue (eWAT). L. Quantification of blots in K. Results are representative of at least two independent experiments. Data are presented as mean ± SEM; ns, not significant; *, P < 0.05; and **, P < 0.01 by Student’s t test (A-J) and one-way ANOVA with Newman-Keuls Multiple Comparison test (L).

We next assessed the effect of PPARγ phosphorylation on glucose homeostasis in HFD-fed mice. Similar to effects observed in chow-fed mice, glucose tolerance was comparable between genotypes (Fig. 1C and Suppl Fig. 2C). In contrast, insulin tolerance testing in HFD-fed mice revealed PPARγ^A/A^ mice to be more responsive to insulin, as evidenced by a significantly enhanced glucose lowering response (Fig. 1D). To better understand the insulin sensitivity phenotype, we performed hyperinsulinemic-euglycemic clamps on HFD-fed mice. We observed increases in the glucose infusion rate (GIR) required to maintain euglycemia in mutant mice compared to controls, indicating improved insulin sensitivity *per se* (Fig. 1E). The primary factor contributing to the increased GIR is elevated whole-body glucose utilization, with pronounced effects observed in skeletal muscle but not white adipose tissue (WAT) (Fig. 1F–1H). Of note, there was no significant difference in the ability of insulin to suppress endogenous glucose production (EGP) (Fig. 1I). These findings suggest that the improved insulin sensitivity is due to greater glucose uptake into tissues and is not due to decreased glucose production. The improved glucose uptake in muscle rather than fat is highly suggestive of tissue cross-talk. Importantly, we observed a three-fold increase in the ability of insulin to suppress non-esterified fatty acid (NEFA) levels in PPARγ^A/A^ mice (Fig. 1J).

We also examined biochemical indicators of insulin sensitivity and insulin signaling. PPARγ^A/A^ mice exhibited a modest but significant increase in insulin-stimulated AKT phosphorylation within skeletal muscle but not liver, inguinal WAT (iWAT), or epididymal WAT (eWAT) (Figs. 1K–1L). These results strongly support the notion that phosphorylation of PPARγ at S273 is sufficient to promote obesity-induced insulin resistance.

### Effect of TZDs on PPARγ^A/A^ mice

The TZD class of anti-diabetic drugs, which are both agonists and prevent phosphorylation of PPARγ S273, are potent insulin sensitizers. However, these drugs also have adverse effects, such as weight gain, fluid retention, and bone loss [22]. Whether these side effects are completely separable from insulin sensitization is a key unanswered question. Multiple small-molecule non-agonist ligands have successfully separated beneficial from adverse effects [17, 23, 24]. To test whether this phenomenon holds true in a genetic model, we assessed PPARγ^A/A^ mice for TZD-associated side effects. First, adipocytes in eWAT of PPARγ^A/A^ mice had similar cell surface area to cells from control mice (Fig. 2A). In addition, PPARγ^A/A^ mice fed HFD showed no decrease in hematocrit, as an indicator of fluid retention (Fig. 2B), compared to controls. Importantly, using microquantitative computed tomography (microCT) we also observed no change in tissue mineral density (TMD) or trabecular bone volume fraction among genotypes (Fig. 2C–2D). Cortical bone parameters, including thickness, mineralization and periosteal perimeter, were similarly unchanged (data not shown). Thus, while mice harboring the S273A knock-in allele recapitulate the PPARγ agonist-like improvements in insulin sensitivity, they avoid the adverse effects typically associated with TZDs. Treatment of HFD-fed PPARγ^A/A^ mice with the full agonist of PPARγ, rosiglitazone, revealed a further improvement in insulin sensitivity (Fig. 2E). These effects included the adverse effects, weight gain and increased adiposity (Fig. 2F-G). These results indicate that while PPARγ^A/A^ mice show an improvement in glucose homeostasis, these animals are still susceptible to certain beneficial effects of TZD treatment and to the full adverse effects of these drugs.

**Figure 2.**
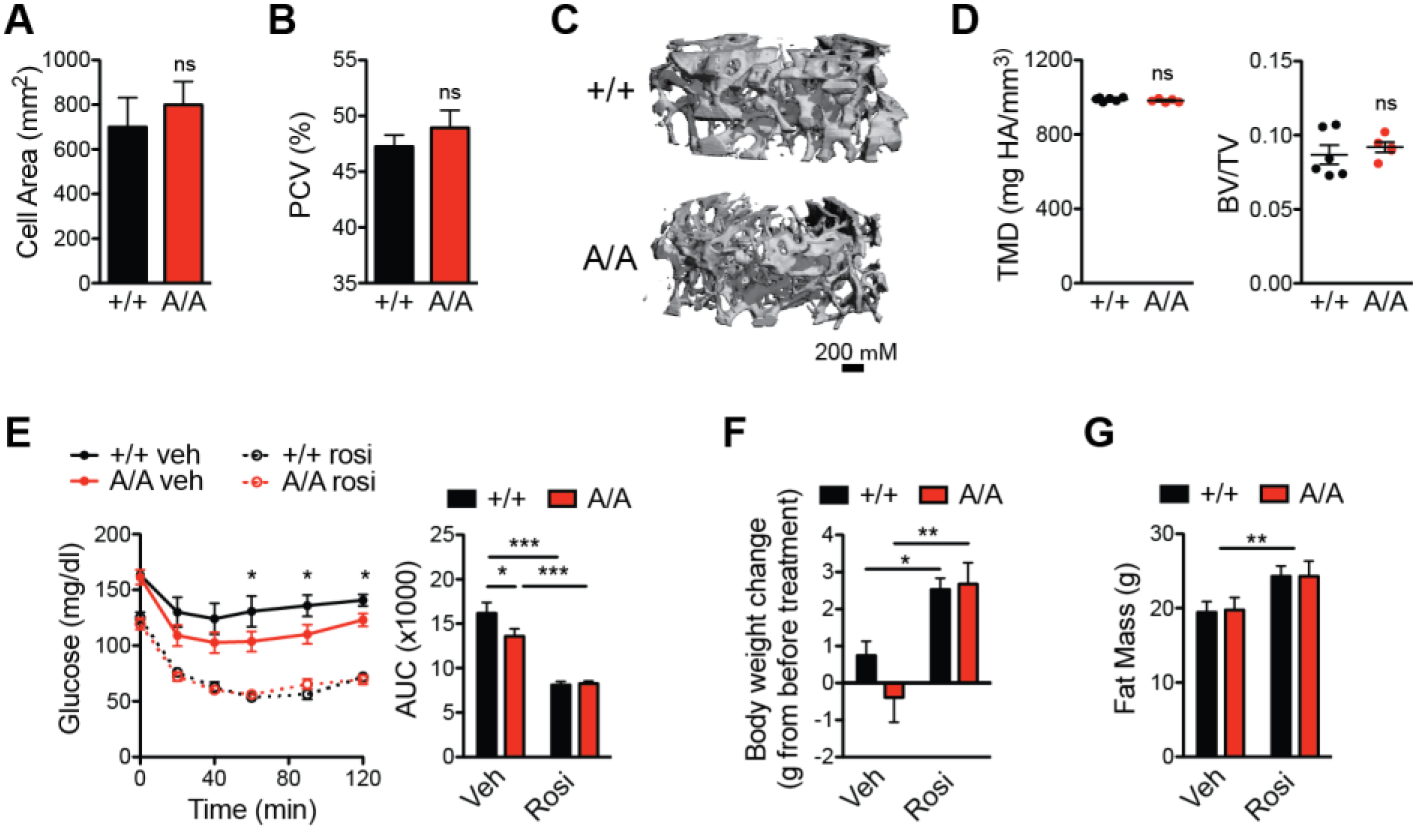
PPARγA/A mice respond to PPARγ agonist treatment. A-D HFD-fed PPARγA/A mice (A/A) do not innately exhibit the adverse effects associated with PPARγ agonism. A. Average cell area of adipocytes from epididymal adipose tissue after HFD feeding (n = 3). B. Hematocrit (packed cell volume, PCV; n = 6 +/+, 5 A/A). C-D. Microquantitative computed tomography (microCT) analysis of trabecular bone from femurs of 6-month old mice maintained on HFD (n = 6 +/+, 5 A/A), C. Representative image of microCT, D. Parameters from microCT analysis, including tissue mineral density (TMD, left) and trabecular bone volume fraction (BV/TV, right). Data are presented as mean ± SEM; no significant differences were observed. E-G. Both wildtype and A/A mice respond to treatment with PPARγ agonist rosiglitazone. After HFD feeding, mice were treated with either vehicle or rosiglitazone for 10 days (8 mg/kg body weight; n = 7 per genotype). E. Insulin tolerance test (1.5 U insulin/kg body weight); right, area under the curve (AUC). F. Body weight gained over course of treatment. G. Total fat mass at end of treatment. Data are presented as mean ± SEM; ns, not significant; *, P < 0.05; **, P < 0.01; and ***, P < 0.001 by one-way ANOVA with Newman-Keuls Multiple Comparison (E-F) and two-way ANOVA (G).

### Expression of Gdf3 is regulated by PPARγS273 phosphorylation

As PPARγ is phosphorylated in obese states and is highly expressed in adipose tissue, we examined transcriptional differences between control and littermate PPARγ^A/A^ eWAT from mice on HFD. Differential gene expression analysis following RNA-sequencing in eWAT revealed one striking result in the mutant animals: a strong and significant (p < 10^-12^) downregulation of Gdf3, a secreted protein and member of the TGF-β superfamily (Fig. 3A). Using qPCR, we confirmed *Gdf3* was expressed at lower levels in both PPARγ^A/A^ iWAT and eWAT from HFD-fed mice when compared to littermate controls (Fig. 3B). Consistent with previously published data, eWAT Gdf3 levels increase in obesity [25]. *Gdf3* expression shows a 35-fold increase in the eWAT of WT mice maintained on a HFD for 25 weeks compared to chow-fed controls. In contrast, the induction of *Gdf3* by HFD was reduced 80 percent in the eWAT of PPARγ^A/A^ mice. No significant difference was observed between genotypes on a standard chow diet (Fig. 3C). We next investigated Gdf3 regulation by non-genetic modulation of PPARγ S273 phosphorylation using small-molecules that can limit S273 phosphorylation of PPARγ. Mice were treated with rosiglitazone, a PPARγ agonist or with MEK inhibitors, GSK1120212 (Trametinib) or PD0325901 to reduce PPARγ phosphorylation. *Gdf3* responses were significant in all pharmacological models tested. *Gdf3* mRNA levels decreased significantly in response to rosiglitazone, Trametinib, and PD0325901 by 60%, 68%, and 50% respectively (Fig. 3D).

**Figure 3.**
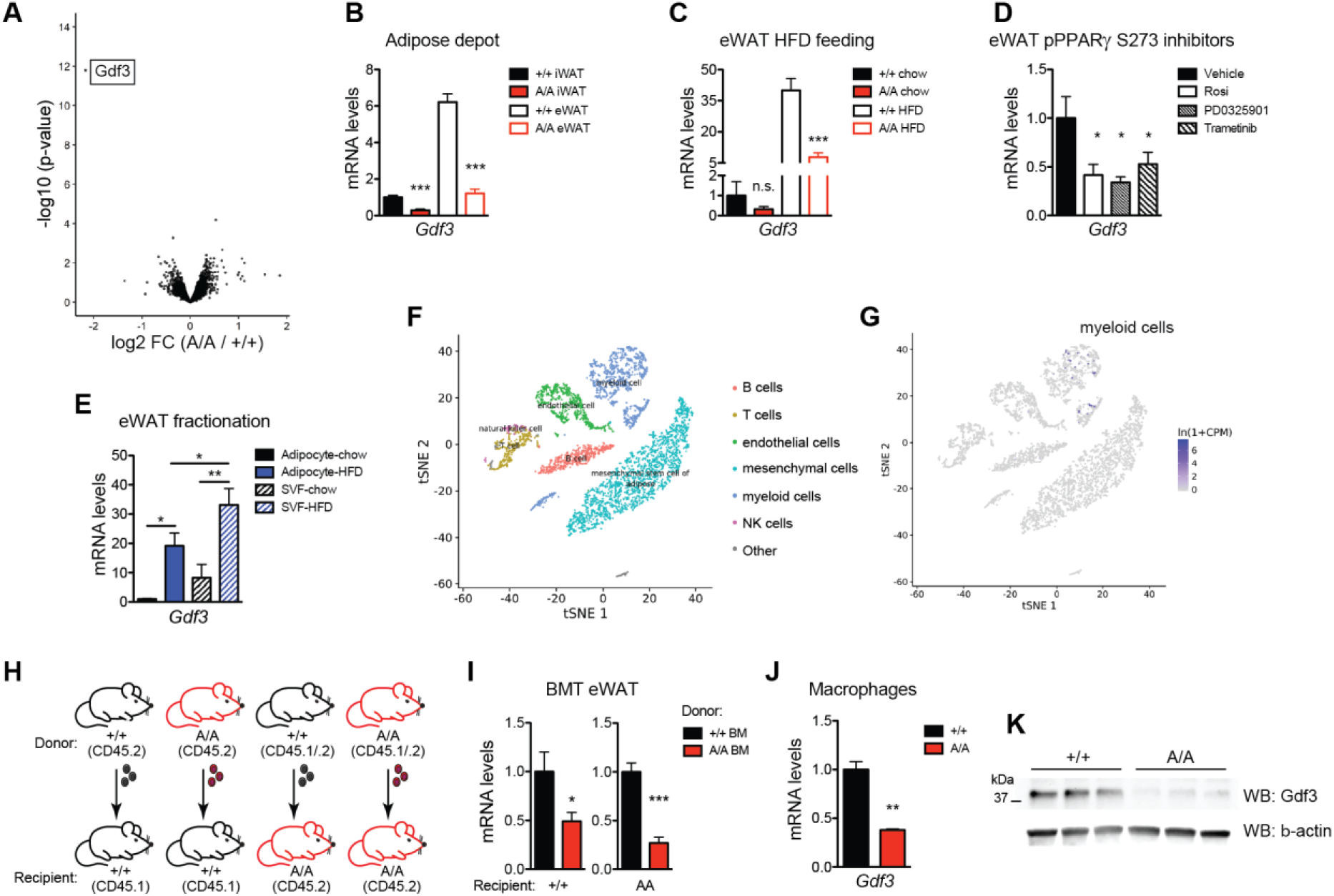
Mice with a non-phosphorylatable allele of PPARγ have decreased *Gdf3* expression in adipose tissue. A. Volcano plot from RNA-seq analysis on epididymal adipose tissue (eWAT) of obese wildtype (+/+) and PPARγA/A (A/A) mice, with differentially expressed gene *Gdf3* highlighted. B. qPCR validation of decreased *Gdf3* mRNA levels in eWAT and inguinal adipose tissue (iWAT) of HFD-fed A/A mice. C. *Gdf3* mRNA levels in eWAT from mice on chow or HFD. D, *Gdf3* mRNA levels in obese adipose tissue also decrease following treatment with known inhibitors of PPARγ S273 phosphorylation (rosiglitazone or either of two MEK inhibitors, PD0325901 or Trametinib; n = 4-ø). E. Fractionated adipose tissue indicates *Gdf3* is highly expressed in both the SVF and adipocyte fractions of eWAT from mice on HFD. F-G. Single Cell RNA-sequencing of SVF from chow-fed wildtype mice shows detectable expression only in the myeloid population H-l. Generation of bone marrow chimeras reveals dominant role of hematopoietic cells in decreased *Gdf3* expression of A/A mice. H. Scheme used for bone marrow transplantation study, including use of congenic C57BL/6 strains carrying functionally equivalent alleles of the pan leukocyte marker CD45. I. *Gdf3* mRNA levels in eWAT after HFD treatment of chimeric mice (n = 7-8). J-K. Thioglyco Hate-elicited peritoneal macrophages from A/A mice exhibit decreased Gdf3 (J) mRNA and (K) protein levels. Data are presented as mean ± SEM; *, P < 0.05; **, P < 0.01; and ***, P < 0.001 by Student’s t test (B-C, per tissue/conditition; l-J) and one-way ANOVA with Newman-Keuls Multiple Comparison test (D-E).

### GDF3 is regulated by PPARγ phosphorylation in myeloid cells

Due to the heterogeneous composition of adipose tissue, we examined whether Gdf3 levels were found within a population of fractionated adipocytes or the stromal vascular fraction (SVF). Although HFD treatment increased *Gdf3* expression in both adipocyte and SVF fractions, higher *Gdf3* mRNA levels were observed in the SVF (Fig. 3E). Single-cell RNA-seq of mouse SVF suggests that Gdf3 is localized to myeloid cells within the SVF [26] (Fig. 3F-G). To assess the contribution of macrophage-specific PPARγ S273 phosphorylation to obesogenic Gdf3 expression, we generated bone marrow chimeras and asked whether obese WT mice with a PPARγ^A/A^ immune system will have altered Gdf3 expression (Fig. 3H). Strikingly, mice with a hematopoietic compartment reconstituted from PPARγ^A/A^ donors exhibited lower *Gdf3* expression in eWAT regardless of recipient genotype (Fig. 3I). Thus, transplantation with PPARγ^A/A^ bone marrow recapitulated the decreased *Gdf3* levels observed with whole-body PPARγ^A/A^ animals. Moreover, peritoneal macrophages from PPARγ^A/A^ mice revealed a 65% reduction in the levels of *Gdf3* mRNA and an 80% reduction in protein levels (Fig. 3G–3H). Together, these data provide strong evidence for the regulation of macrophage Gdf3 by PPARγ S273 phosphorylation.

To determine whether the differential levels of Gdf3 levels in adipose tissue were due to decreased numbers of Gdf3 expressing cells or to specific modulation of gene expression, we performed detailed immunophenotyping on eWAT from wildtype and PPARγ^A/A^ mice maintained on HFD for 16 weeks. None of the macrophage or T cell populations examined showed significant differences when either the cell number or the percentage of gated cells was examined between genotypes. Macrophage populations were gated according to the strategy outlined in Suppl. Fig. 5A. No differences were observed in total macrophage number or percent of gated live, hematopoietic cells (CD45^+^) that were not eosinophils (Siglec F^-^) (Suppl. Fig. 5B). Nor were differences in macrophage percent of cells expressing an M1 marker (CD11c^+^) or an M2 marker (CD301^+^) (Suppl. Fig. 5C). PPARγ is highly expressed in T regulatory cells (Treg), a cell type which can control tissue inflammation including macrophage recruitment [8, 27]. Treg populations were gated according to the strategy outlined in Suppl. Fig. 5D. No differences were observed in total Treg number or percent of gated live, hematopoietic cells (CD45^+^) that expressed the T cell receptor, CD4 co-receptor, and transcription factor Foxp3 (TCRb^+^ CD4^+^ Foxp3^+^) (Suppl. Fig. 5E). Similarly, tissue-resident Tregs (KLRG1^+^ ST2^+^), had no significant differences (Suppl. Fig. 5F). Coupled with the lack of cell type specific gene regulation, these data support the conclusion that similar levels of hematopoietic cells are present in PPARγ^A/A^ adipose tissue. These findings strongly suggest the differences in Gdf3 expression levels in PPARγ^A/A^ adipose tissue reflect specific transcriptional control by PPARγ S273 phosphorylation.

### Gdf3 limits BMP signaling

Members of the TGF-β superfamily, including BMPs, GDFs, and TGF-β proteins can affect insulin sensitivity, cellular differentiation, and obesity [28–31]. These proteins typically work either through BMP-type receptors or TGF-β–type receptors. Relatively little is known of mammalian Gdf3 function on glucose homeostasis *in vivo*. Two seemingly incompatible models have been proposed to explain how extracellular Gdf3 signaling is propagated. One model suggests that Gdf3 can inhibit BMP-receptor signaling, possibly by competing with other ligands for receptor binding [32, 33]. In the other model, Gdf3 *activates* TGF-β type receptors (ALK7/Cripto), especially at super-physiological concentrations of 300-500 ng/ml [34–36]. We examined whether Gdf3 can affect either signaling system using a WT full length Gdf3 cDNA expressed in human HEK293 cells. Activin, a TGF-β receptor ligand activates Smad2/3 phosphorylation. In contrast, Gdf3 had little impact on TGF-β signaling. Activin neither promotes nor inhibits BMP-receptor signaling as measured by phosphorylation of SMAD1/5/9. However, cells expressing Gdf3 demonstrate a strong reduction of SMAD1/5/9 phosphorylation (Fig. 4A).

**Figure 4.**
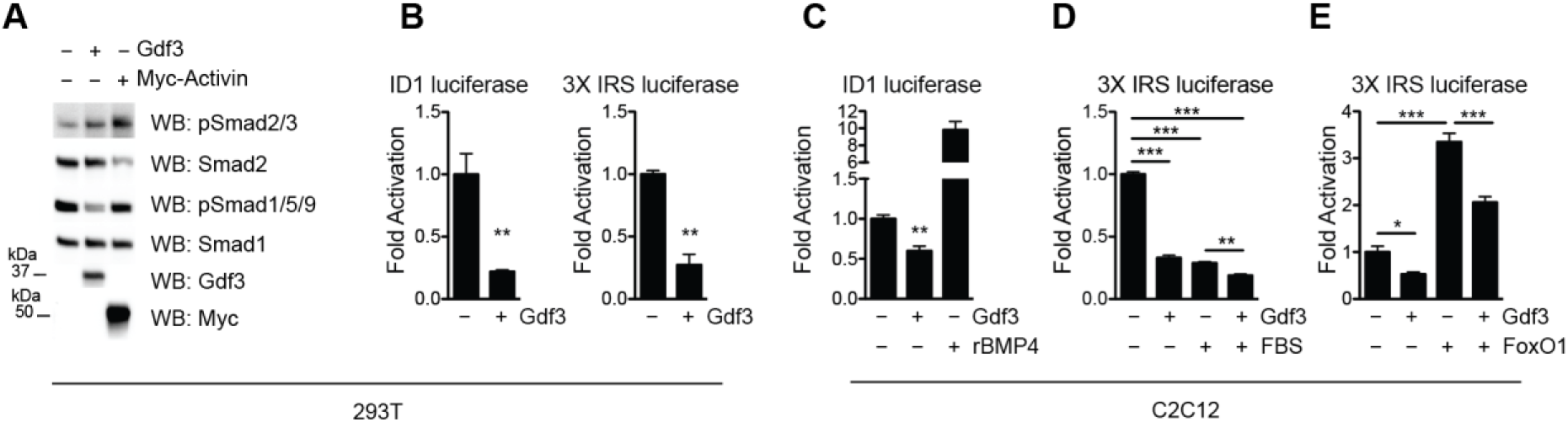
Gdf3 suppresses BMP and FoxO signaling. A. Immunoblot analysis of HEK293 cells transfected with Gdf3 or Myc-Activin constructs. Cells were deprived of serum for 16 h prior to whole-cell extract. B. Luciferase assay using a BMP-responsive element (ID1 luciferase) ora FoxOl-responsive element (3X IRS) driving luciferase reporters in serum-starved HEK293 cells. C. ID1 luciferase assay in C2C12 myoblasts that have been deprived of serum. Overnight treatment of 100 ng/mL rBMP4 demonstrates reporter activation. D-E. 3X IRS luciferase assay in C2C12 myoblasts that are (D) in the presence (+ FBS) or absence of serum or (E) activated by a co-transfected FoxOl construct. Data are presented as relative activity after normalization to Renilla luciferase and shown as mean ± SEM; *, P < 0.05; “, P < 0.01; and ***, P < 0.001 by Student’s t test (B-C) and one-way ANOVA with Newman-Keuls Multiple Comparison test (D-E).

To understand the integrated effects of Gdf3 over time, we performed luciferase reporter assays to quantify Gdf3 transcriptional consequences. In reporter assays, BMP signaling promotes SMAD 1/5/9 binding to the Id1-Luc promoter. We observed an 80% reduction of luciferase activity with Gdf3 overexpression compared to control cells expressing an empty vector (Fig. 4B). In some contexts, BMP signaling may affect the activation of Akt, a key component of insulin signaling [37]. Accordingly, we further examined the impact of Gdf3 on Akt signaling pathways using reporter assays. Unlike SMAD proteins, Akt lacks direct transcriptional effects. However, Akt kinase activity rapidly triggers the nuclear exclusion of transcription factor FoxO1 and prevents binding to a multimerized 3x IRS-Luc reporter [38]. Consistent with increased AKT phosphorylation, overexpression of Gdf3 decreased FoxO1 transcriptional activation (Fig. 4B), supporting a model of elevated Akt activity.

We further confirmed these results in a second cell type, C2C12 myoblasts. As in HEK293 cells, Gdf3 decreases Id1-Luc by 50% compared to control cells. As a positive control, BMP4 activates BMP receptor signaling and stimulates Smad1/5/9 activity (Fig. 4C). To better understand the regulation of Akt by Gdf3, we again utilized the FoxO1 3x IRS-Luc reporter. To increase FoxO1 transcriptional activity, cells were deprived of fetal bovine serum to increase FoxO1 nuclear occupancy. Gdf3 suppressed FoxO1 transcriptional activity by 70%. A further decrease in FoxO1 reporter activity was observed in Gdf3 expressing cells treated with FBS (Fig. 4D). Gdf3 inhibits FoxO1 reporter activity both with endogenous levels of FoxO1 and in the presence of a FoxO1 expression plasmid. These data suggest that Gdf3 is a BMP inhibitor in a class similar to Noggin or Gremlin [39, 40].

### Gdf3 affects glucose uptake in vitro

Based on the increased skeletal muscle glucose uptake in HFD fed PPARγ^A/A^ mice, we wanted to determine if regulating Gdf3 levels in muscle cells could affect insulin-dependent glucose uptake. To test this, we transduced differentiated C2C12 myotubes with either a GFP control or Gdf3 adeno-associated virus (AAV) expressing GFP (AAV2/DJ-H2B-GFP), or an AAV expressing Gdf3 (AAV2/DJ-CAG-Gdf3) (Fig. 5A). Gdf3 overexpression in these cells led to decreased glucose uptake in the presence of insulin, compared to GFP expressing cells. No differences between groups were observed under baseline (no insulin), (Fig. 5B).

**Figure 5.**
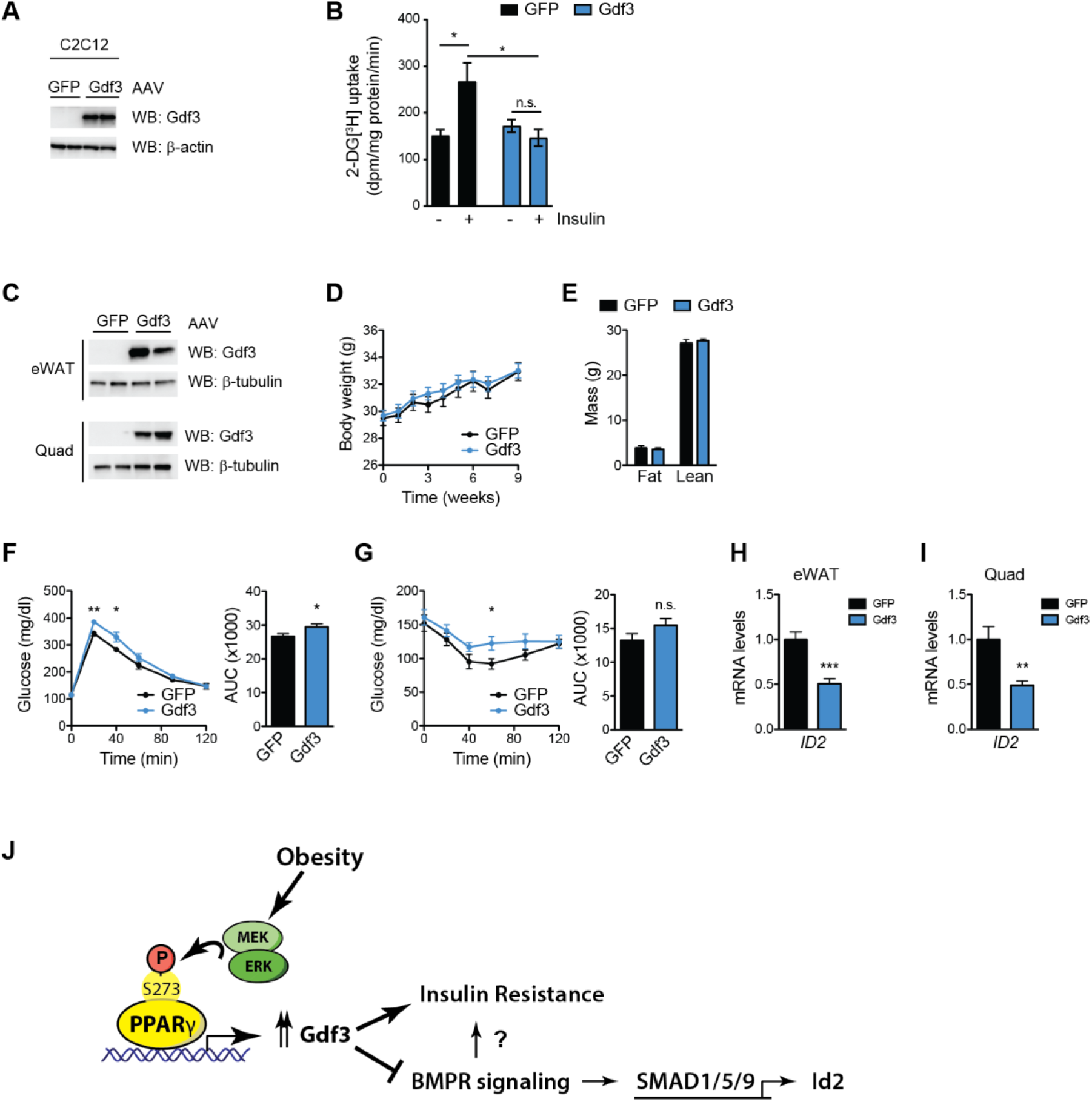
Gdf3 promotes insulin resistance. A, B. C2C12 myotubes transduced with AAV-GFP orAAV-Gdf3 and analyzed by immunoblot for Gdf3 and β-actin (A), [3H] 2-deoxy-D-glucose (2-DG) uptake assays, n = 6 wells per group (B). C-l. As a gain-of function model, mice were administered AAV-GFP orAAV-Gdf3, n = 10 mice per group. Representative immunoblots for Gdf3 and β-tubulin from eWAT and quad from two mice from each group (C). Body weights of mice; week Ũ represents the values on the day of AAV injections (D). Body composition at 6 weeks post injections (E). Insulin tolerance test (0.75 U insulin/kg body weight) 3 weeks post injections (F). Glucose tolerance test (2 g glucose/kg body weight) 4 weeks post injections (G). Right, area under the curve (AUC)forthe respective test (F-G). Relative Id2 mRNA levels from eWAT (H) and quadriceps (I). L. Model illustrating novel role for PPAR⌧ S273 phosphorylation and Gdf3 in impaired insulin action with obesity. Data are presented as mean ± SEM; *, P < 0.05; **, P < 0.01; and ***, P < 0.001 by two-way ANOVA with Sidak’s Multiple Comparison test (B) and Student’s t test (C-I).

### Gdf3 is sufficient to induce insulin resistance in vivo

To investigate the effect of Gdf3 *in vivo,* we administered AAV-GFP or Gdf3 into the visceral fat pads (eWAT) and muscle (quadriceps and gastrocnemius) of lean WT mice resulting in a detectable increase in protein levels of Gdf3 (Fig. 5C). Forced Gdf3 expression did not affect body weight, body composition, energy expenditure, or food intake of mice fed a chow diet (Fig. 5D, 5E and data not shown). Importantly, mice overexpressing Gdf3 had impairments in both insulin tolerance and glucose tolerance tests (Fig. 5F and 5G). These data indicate that increased levels of Gdf3 in these tissues are sufficient to negatively impact glucose homeostasis, independently of changes in body weight. The inhibitors of differentiation (Id) genes, including Id2 are canonical targets of BMPR signaling [41]. We examined the effect of AAV Gdf3 expression on *Id2* mRNA levels in eWAT and quadriceps. In both tissues, we observed significant downregulation (Fig 5H and 5I, consistent with inhibition of BMPR signaling *in vivo.* These effects of Gdf3 to promote impaired glucose homeostasis in healthy, lean mice suggest that the elevated Gdf3 levels observed in obesity may contribute to insulin resistance.

## Discussion

The link between PPARγ biology and insulin sensitivity is both robust and complex. Patients treated with TZD PPARγ agonists experience durable reduced glycemia and insulin sensitivity. These effects were long thought to depend on PPARγ-driven adipogenesis—the generation of new, healthy adipocytes from precursor cells. The properties of PPARγ agonists to promote adipocyte differentiation cause weight gain, but also decrease bone formation leading to increased risk of fractures. The demonstration that blocking PPARγ S273 phosphorylation can improve insulin sensitivity independently of receptor agonism or effects on bone, suggest new therapeutic approaches to insulin sensitizers.

Increased PPARγ S273 phosphorylation induced by ERK kinase in obesity contributes to insulin resistance. Our study shows that preventing S273 phosphorylation genetically is sufficient to maintain insulin sensitivity in obesity, as manifested by improved insulin tolerance and increased whole body glucose utilization (Fig. 1). Consistent with this interpretation, in lean wild type chow-fed mice in which the receptor is not phosphorylated, we do not observe differences in glucose homeostasis from littermate PPARγ^A/A^ mice. (Suppl. Fig. 2). This suggests that the observed phenotypic changes are due to the contribution of obesity, rather than developmental effects. Moreover, as PPARγ S273 phosphorylation is independent of receptor agonism, we do not observe TZD-like side effects in PPARγ^A/A^ mice (Fig. 2).

We identify Gdf3 as a transcriptional target of PPARγ S273 phosphorylation in obese adipose tissue. Gdf3 is expressed in adipose tissue in both the SVF and adipocyte fractions, with highest levels observed in macrophages (Fig. 3). PPARγ S273 phosphorylation activates Gdf3 transcription; accordingly, studies of macrophage-specific PPARγ deficiency identified strongly reduced Gdf3 mRNA levels, along with many other genes [42]. PPARγ expression in macrophage is required for the regulation of multiple transcriptional programs and for full TZD responses. Gdf3 was the most strongly downregulated gene in expression profiling experiments of macrophage-specific PPARγ deletion [42]. While macrophage PPARγ-deficient mice express low Gdf3 levels which would predict insulin sensitivity, they nonetheless exhibit insulin resistance due to increased secretion of multiple pro-inflammatory cytokines [7, 42–45]. In contrast, PPARγ^A/A^ mice do not have significant differences in macrophage number, relative polarization, or cytokine expression levels (Supp. Fig. 5).

Relatively little is known of mammalian Gdf3 function on glucose homeostasis *in vivo.* Gdf3 is highly expressed in white adipose tissue with levels increasing with age and obesity [25, 36, 46]. Our data support a model in which Gdf3 blocks beneficial BMP receptor signaling, decreasing phosphorylation of SMAD 1/5/9, and inhibiting Id2, a SMAD target gene (Fig. 4 and 5).

Our work is consistent with studies demonstrating improvements in glucose homeostasis by increasing levels of BMP4. Conversely, BMP4 KO mice exhibit impaired glucose tolerance [47, 48]. Heterozygous deletion of the type 1 BMP receptor, *Bmpr1a* causes impaired glucose tolerance [49], while adipocyte-specific receptor deletion does not [50]. Similar salutary effects of glucose homeostasis have been observed in mice treated with BMP6, BMP7, and BMP9 [28, 30, 31]. BMPs can drive cellular differentiation including potentiation of brown and beige fat activation [47, 48, 51]. In this context, tight control of BMP signaling is maintained by inhibitory proteins including Gdf3, Noggin and Gremlin [33, 52].

An emerging literature is developing on the contribution of macrophages to catecholamine production and degradation. Macrophages expressing Gdf3 have been implicated in controlling catecholamine degradation and rates of lipolysis [46]. Mice with global deletion of Gdf3 from birth maintain lean body masses when challenged with HFD [34, 53]. Conversely, Gdf3 has been reported to induce severe and persistent obesity in an adenovirus overexpression model [25]. Our results are different from these earlier findings. We observe no differences in energy balance in our models of decreased Gdf3 levels with the PPARγ^A/A^ mice or with exogenous Gdf3 expression using AAV (Fig. 5). These differences may be due to the role of Gdf3 in embryonic development [33, 35] versus the late timing of Gdf3 expression changes driven by differential obesity-mediated phosphorylation of S273 PPARγ.

Future studies will determine whether Gdf3 inhibits BMP-receptor signaling by binding to the BMP receptor or through inhibitory binding to other extracellular BMP receptor ligands. We find that Gdf3 is a factor which mediates the adverse consequences of obesity-mediated PPARγ S273 phosphorylation. Further study into the mechanisms of Gdf3 action will be beneficial to understand whether this is an appropriate future therapeutic target.

## Materials and Methods

### Generation of PPARγ^S273A^ mice

We obtained a C57BL/6 bacterial artificial chromosome (BAC) containing the *Ppar* locus (Invitrogen/RPCI). An 8.9 kb fragment preceding and including exon 5 was amplified by PCR (Phusion DNA Polymerase, NEB) and cloned into a plasmid encoding both a floxed neomycin resistance cassette and a diphtheria toxin negative selection marker. Site-directed mutagenesis was used to alter the codon for serine 273 to alanine (TCA to GCA). Lastly, a 1.6 kb 3’ homology arm was similarly amplified and cloned into the plasmid. C57BL/6 ES cells were electroporated with the linearized targeting construct. Using positive and negative selection, 8 ES cell clones were isolated and two were injected into Balb/c blastocysts. The resulting progeny gave rise to highly chimeric offspring. These chimeras were mated to C57BL/6J mice to obtain germline transmission of the mutant allele. To remove the floxed neomycin cassette, these germline offspring were crossed to mice expressing Cre in the female germline (Zp3-Cre). The progeny from these crosses were found to have deleted the neomycin cassette successfully. These heterozygous mice (PPARγ^S273A/+^) were mated to generate homozygous knock-in mice PPARγ^S273A/S273A^ or PPARγ^A/A^. Mice were backcrossed onto C57BL/6J for 6 generations, after which mice heterozygous for the PPARγ^S273A^ allele (PPARγ^S273A/+^) were bred to generate experimental cohorts. The PPARγ^S273A^ allele was detected by PCR genotyping with primers 5’-CAGGAGGCTGAGCAGGTGTGTT-3’, 5’-TCCAGACTGCCTTGGGAAAA-3’, and 5’-AGCACACATGTACCCAACAT-3’ (34 cycles, 96°C, 30s; 65°C, 45s; 72°C, 45s), where the wildtype and knock-in alleles produced PCR products of 319 bp and 217 bp, respectively. To minimize phenotypic variability from a mixed 6J/6N background, PPARγ^S273A/+^ mice were genotyped for single nucleotide polymorphisms (SNPs) that distinguish between C57BL6/J and C57BL6/N alleles [54]; mice homozygous for at least 9 SNPs specific to the C57BL/6J substrain on distinct chromosomes, including the 6/J null allele encoding nicotinamide nucleotide transhydrogenase *(Nnt)* [55], were used to establish cohorts for subsequent experiments.

### Animal experiments

Animal experiments were performed with approval from the Institutional Animal Care and Use Committees of The Harvard Center for Comparative Medicine, Brigham and Women’s Hospital, and Beth Israel Deaconess Medical Center. Mice were fed a standard chow diet (13% kcal fat, LabDiet, no. 5053) or a high-fat diet (60% kcal fat, Research Diets, no. D12492i) for the indicated durations. For the *in vivo* AAV experiments, adult male C57BL6/J mice were purchased from Jackson Labs and allowed to acclimate to our animal facility for 1 week. Each AAV was injected bilaterally into the quadriceps and gastrocnemius muscles at a dose of 1.5 x 10^8^ vg in 30 μL of PBS per injection and into the epididymal fat depots at 2.5 x 10^8^ vg in 50 μL of PBS per injection. These viral injections were performed under isoflurane anesthesia and mice were given Buprenorphine SR Lab as the perioperative and postoperative analgesic prior to surgery at a dose of 0.5 mg/kg. Glucose and insulin tolerance tests were performed on mice fasted for 16 and 4 hours, respectively. Following an initial blood glucose measurement post-fast, tolerance to an intraperitoneal (IP) injection of glucose (1 g/kg) or insulin (1.5-2 U/kg) was assessed by change in blood glucose over a period of 120 minutes. Glycemia was measured by tail vein bleeds at the indicated times using a Contour Next EZ glucometer (Bayer). Hyperinsulinemic-euglycemic clamp experiments were performed as previously described [19]. For PPARγ ligand treatment prior to ITT, diet-induced obese mice were IP injected daily for 10 days with rosiglitazone (Cayman Chemical; 8 mg/kg) before insulin tolerance testing. Rosiglitazone was dissolved in dimethylsulphoxide (DMSO) and diluted into saline with 2% Tween-80 for injection. Administration of MEK inhibitors Trametinib and PD0325901 (Selleckchem) were as previously described [19]. Body composition measurements were assessed with an EchoMRI 3-in-1 (Echo Medical Systems) on conscious mice. For biochemical analysis of insulin signaling, obese mice of both genotypes were fasted for 4 hours prior to IP injection of insulin (10 U/kg). Ten minutes after insulin injection, white adipose tissue, liver, and gastrocnemius were collected for subsequent protein analysis. At the end of experiments, tissues were snap-frozen in liquid nitrogen and stored at −80°C until processing. Alternatively, samples of adipose tissue were fixed in formaldehyde, embedded in paraffin, and stained with H&E for determination of adipocyte area using ImageJ software, as previously described [56]. For measures of hematocrit, a small volume of whole blood was centrifuged in a microhematocrit capillary tube, and the height of the column of packed cells was expressed as a percentage of the height of the total blood column (visibly hemolyzed samples were excluded).

### Indirect calorimetry

For measurements of metabolic rate and food intake, we placed mice within the CLAMS (Columbus Instruments) indirect calorimeter. One week prior to monitoring, mice were surgically implanted with wireless body temperature probes (Starr Scientific). Mice were acclimated for 24 h followed by 24 h measurement of VO_2_, VCO_2_, RER, locomotor activity, food intake at 23°C while on a HFD. Energy expenditure was calculated with the Weir equation [57]. Calories consumed were calculated by multiplying hourly food intake by the 5.21 kcal/g caloric value of the 60% HFD. Energy balance was calculated by subtracting hourly food intake from hourly energy expenditure. Analysis with ANOVA and ANCOVA were performed using CalR [58].

### Microquantitative computed tomography

For microCT analysis, a Scanco Medical μCT 35 system with an isotropic voxel size of 7 μm was used to image the femur. Scans were conducted in 70% ethanol using an X-ray tube potential of 55 kVp, an X-ray intensity of 0.145 mA and an integration time of 600 ms. Analysis was performed according to published guidelines [59]. From the scans of the femur, a region beginning 0.28 mm proximal to the growth plate and extending 1mm proximally was selected for trabecular bone analysis. A second region 0.6 mm in length and centered at the midpoint of the femur was used to calculate diaphyseal parameters. A semi-automated contouring approach was used to distinguish cortical and trabecular bone. The region of interest was thresholded using a global threshold that set the bone/marrow cut-off at 452.65 mg HA/cm^3^ for trabecular bone and 579.86 mg HA/cm^3^ for cortical bone. 3-D microstructural properties of bone, including TMD and bone volume fraction (BV/TV), were calculated using software supplied by the manufacturer.

### RNA isolation and gene expression analysis by qPCR, RNA-seq, or scRNA-seq

RNA Isolation: For experiments utilizing RNA-seq or qPCR, total RNA was isolated from cells or tissues using TRIzol reagent (Thermo Fisher) followed by a Direct-zol RNA MiniPrep kit (Zymo Research) according to the manufacturer’s instructions. The RNA was reverse-transcribed using Multiscribe Reverse Transcriptase and a high-capacity cDNA reverse transcription kit (Applied Biosystems).

Quantitative real-time PCR (qPCR) was performed using synthesized cDNA with SYBR Select Master Mix (Applied Biosystemes) on a Light Cycler 480 II (Roche) or the QuantStudio 5 RealTime PCR System (Applied Biosystems). Relative mRNA expression was determined by the △△-Ct method normalized to TATA-binding protein (TBP) levels. The list of primers used in this study is included in Supplementary Table S1.

RNA-sequencing (RNA-seq): 400ng of total RNA was used as input for RNA-seq Library construction, which consisted of rRNA removal via the NEBNext rRNA depletion kit (E6310X) and conversion to cDNA using the NEBNext Ultra II Non-Directional RNA Second Strand Synthesis Module (E6111L), both per manufacturer’s instructions. cDNA was diluted 1:15 and subsequently tagmented and amplified for 12 cycles using the Nextera XT DNA Library Preparation Kit (Illumina FC-131). Individual libraries were QC’d by Qubit and Agilent Bioanalyzer, pooled, and sequenced at a final loading concentration of 1.8 pM on a NextSeq500 (36bp x 36bp paired-end reads). Sequencing reads were demultiplexed using bcl2fastq (v2.20.0) and aligned to the mm10 mouse genome by with HISAT2 (v. 2.0.5 [60]. PCR duplicates and low-quality reads were filtered by Picard (https://broadinstitute.github.io/picard). Filtered reads were assigned to the annotated transcriptome and quantified using featureCounts [61]. Normalization and differential expression analysis were performed using EdgeR (v 2.9.2) [62].

Single-cell RNA-seq (scRNA-seq): Adipose tissue SVF scRNA-seq data were accessed from the Tabula Muris data set. These data were from the combined subcutaneous adipose tissue, gonadal adipose tissue, mesenteric adipose tissue, and interscapular brown adipose tissue, from 4 male and 3 female, chow-fed 10-15 week old C57BL/6JN mice obtained from Charles River Labs [26].

### Preparation of cell or tissue lysates and immunoblotting

Whole-cell extracts were prepared in RIPA lysis buffer containing 50 mM Tris pH 7.5, 150 mM NaCl, 1% NP-40, 0.5% sodium deoxycholate, 0.1% SDS, and 1× HALT protease and phosphatase inhibitor cocktail (Thermo Fisher). For tissue lysates, frozen tissue was pulverized in liquid nitrogen followed by homogenization in lysis buffer using stainless steel ball bearings and a TissueLyser (Qiagen). Protein concentration was determined by Pierce BCA protein assay (Thermo Fisher) and lysates separated by SDS-PAGE with Mini-PROTEAN TGX gels (Bio-Rad), transferred to PVDF membranes using the Trans-Blot Turbo transfer system (Bio-Rad), and blotted according to manufacturer’s recommendations for the indicated antibodies. Primary antibodies used in this study include: rabbit anti-phospho-Akt Ser473 (Cell Signaling, 4060S), rabbit anti-Akt (Cell Signaling, 9272S), rabbit anti-β-Tubulin (Cell Signaling, 2146), rat anti-Gdf3 (R&D Systems, MAB953), mouse anti-β-Actin (Sigma, A1978), rabbit anti-phospho-Smad2 Ser465/467/ Smad3 Ser423/425 (Cell Signaling, 8828S), rabbit anti-Smad2 (Cell Signaling, 5339S), rabbit anti-phospho-Smad1 Ser463/465/ Smad5 Ser463/465/ Smad9 Ser465/467 (Cell Signaling, 13820S), rabbit anti-Smad1 (Cell Signaling, 6944S), rabbit anti-Myc-Tag (Cell Signaling, 2278S), rabbit anti-phospho-PPARγ Ser273 (as previously described [14]), and mouse anti-PPARγ (Santa Cruz, sc-7273). Horseradish peroxidase-conjugated secondary antibodies were obtained from Cell Signaling Technologies (anti-rabbit IgG, 7074S; anti-mouse IgG, 7076S, and anti-rat IgG, 7077S). Chemiluminescence assays were performed using SuperSignal West Dura extended duration substrates (Thermo Fisher) and detected with a ChemiDoc XRS+ or ChemiDoc Touch Imaging System (Bio-Rad). Image analyses were performed using Image Lab software (Bio-Rad).

### DNA constructs and AAVs

A non-tagged construct of mouse Gdf3 was generated using the commercial vector pCMV6-Gdf3-Myc-Flag (OriGene Technologies, MR222967), containing a Myc-Flag-tagged Gdf3, and PCR-based mutagenesis to introduce a STOP codon upstream of the C-terminal Myc-Flag tag. The ID1 luciferase reporter was provided by C. Brown. The 3X IRS luciferase reporter was a generous gift from D. Accili. The FoxO1 constructs have been previously described [63]. A Myc-Flag-tagged mouse Activin expression plasmid was purchased from Origene Technologies (MR225191). AAV Gdf3 was generated by subcloning non-tagged Gdf3 into the AAV2/1 vector backbone of AAV-CAG-GFP, a gift from Karel Svoboda via Addgene plasmid 28014 [64]. Viral packaging into the AAV-DJ serotype and titer determination were performed by the Boston Children’s Hospital Viral Core. Premade control AAV2/DJ-CAG-H2BGFP virus (AAV-GFP) was purchased from the same core facility.

### Cell culture and treatment

HEK293 cells were cultured and maintained in DMEM containing 10% heat inactivated FBS (iFBS). C2C12 myoblasts were cultured and maintained in DMEM containing 20% iFBS and were transfected without myogenic differentiation. Transfections were performed with Lipofectamine 2000 and 3000 transfection reagents (Invitrogen), respectively, according to the manufacturer’s recommendations. Adipocyte differentiation was induced by treating 2-days postconfluent cells (either 3T3-L1 preadipocytes or adipose tissue SVF) with 5 μM dexamethasone, 250 μM isobutylmethylxanthine, 1 μM rosiglitazone, and 900 nM insulin in DMEM with 10% iFBS for 48 h. Cells were maintained in culture medium containing iFBS, insulin, and rosiglitazone for another 2 days. Thereafter, cells were cultured in medium containing iFBS only. Lipid accumulation in adipocytes was detected by Oil Red O staining.

### Glucose uptake assays

Confluent C2C12 myoblast cells were induced to differentiate with serum withdrawal to 2% horse serum in collagen coated 6 well or 48 well plates. Differentiated C2C12 myotubes were returned to DMEM containing 20% iFBS and transduced with either AAV-GFP or AAV-Gdf3 (1.2 x 10^11^ vg per well and 1.2 x 10^11^ vg per well in the 6 and 48 well plates respectively). Approximately 36 hours post transduction, the cells from the 6 well plates were washed twice in ice cold PBS and lysed with RIPA lysis buffer for western blot analysis as described above, while the 48 well plates were used for glucose uptake assays using the protocol described in [65] with minor modifications. Briefly, cells were washed twice with serum-free DMEM and incubated in serum-free DMEM for 3 h at 37°C. Cells were then washed 3 times with Krebs-Ringer-HEPES (KRH) buffer (50 mM HEPES pH 7.4, 137 mM NaCl, 1.25 mM CaCl2, 4.7 mM KCl, 5.0 mM, 1.25 mM MgSO4). Cells were incubated in KRH for 30 min then incubated in KRH with or without 200 nM insulin for 20 min before the addition of a deoxy-D-glucose, 2-[1,2-3H (N)]/2-deoxy-D-glucose solution, yielding final assay concentrations of 1 μCi of [^3^H] 2-deoxy-D-glucose and 100 μM 2-deoxy-D-glucose per well. After 5 min the KRH buffer was removed immediately by aspiration. Cells were washed 3 times in ice-cold PBS to inhibit further glucose uptake and wash away unincorporated radiolabeled [^3^H] 2-deoxy-D-glucose. Cells were solubilized with 0.5% SDS. Half of the lysates from each well were used to assay [^3^H] disintegration levels by liquid scintillation counting and the other half was used for normalization to protein concentrations.

### Luciferase reporter assays

HEK293 or C2C12 cells were transfected with a firefly luciferase reporter construct and a fixed amount of DNA, as normalized with empty vector plasmid. Luciferase activity was measured 24-48 h after transfection using the Dual-Luciferase Reporter Assay system (Promega), with quantification of luminescence on a FLUOstar OPTIMA plate reader (BMG Labtech). Cotransfection of CMV-driven *Renilla* luciferase vector was used to normalize for transfection efficiency. Where indicated, cells were rinsed with PBS and switched to DMEM lacking serum prior to luciferase measurement. As a positive control for BMP signaling, recombinant protein of mouse BMP4 (R&D Systems) was added to the medium 16 h before analysis.

### Isolation of macrophages

Peritoneal macrophages were harvested from control and PPARγ^A/A^ mice 4 days after i.p. injection of sterile 3% thioglycolate solution (3 mL per mouse) as previously described [66]. Following collection of peritoneal cells, erythrocytes were lysed using ACK lysis buffer (Gibco), and the remaining cells seeded on dishes in DMEM containing 10% iFBS and allowed to adhere for 2 h. Nonadherent cells were removed, and adherent cells cultured and used as peritoneal macrophages in the experiments.

### Isolation of cells from SVF

The stromal vascular fraction (SVF) containing preadipocytes and mononuclear cells was extracted from adipose tissue. For isolation of preadipocytes, inguinal adipose tissue was digested in a PBS buffer containing 10 mg/mL collagenase D (Roche), 2.4 U/mL dispase II, and 10 mM CaCl2 for 45 min shaking at 37°C. Digestion was neutralized with culture medium consisting of DMEM/F12 supplemented with 10% FBS and penicillin/streptomycin. Cell suspensions were filtered through 100 μm and 40 μm cell strainers, and the SVF cells resuspended in growth medium for expansion. For isolation of mononuclear cells and subsequent immunophenotyping, epididymal and inguinal adipose tissues were digested for 20-30 min at 37°C with collagenase type II (Sigma). Cell suspensions were filtered through a sieve, and the SVF fraction was collected after red blood cell lysis (ACK Lysing Buffer, Gibco) and filtration through a 40 μm cell strainer.

### Immunophenotyping and flow cytometry

Adipose tissue SVF cells were resuspended in FACS buffer (phenol red-free DMEM supplemented with 10 mM HEPES and 2% FCS) and incubated for 30-45 min at 4°C with Fc block and the indicated fluorescence-labeled antibodies for surface molecule staining. For myeloid cell analysis, cells were stained with anti-CD45.2 (clone 104), -CD45.1 (A20), -CD11b (M1/70), -F4/80 (BM8), and -CD301 (LOM-14) antibodies from BioLegend; -CD11c (N418) antibody from eBioscience; and -SiglecF (E50-2440) antibody from BD Biosciences. For lymphoid cell analysis, cells were stained with anti-CD45.2, -CD45.1, -CD19 (6D5), -TCR-β (H57-597), and -CD8 (53-6.7) antibodies from BioLegend; -CD4 (RM4-5), -KLRG1 (2F1), and -IL33R/ST2 (RMST2-2) antibodies from eBioscience; and were fixed, permeabilized and intracellularly stained for Foxp3 (FJK-16s) according to the manufacturer’s instructions (eBioscience). Dead cells were excluded by staining with a fixable LIVE/DEAD dye (ThermoFisher). Cells were analyzed using an LSRII instrument (BD Bioscience) and FlowJo software.

### Bone marrow transplantation

Bone marrow progenitor cells were isolated by flushing femurs and tibiae of donor animals with PBS, and T-cells depleted from BM by magnetic beads. Recipient mice were irradiated with 1000 rads prior to reconstitution with 3-5 × 10^6^ donor cells by retro-orbital (r.o.) injection. Irradiated mice were kept in sterile cages with standard chow diet and antibiotic-treated water for 2 weeks. After 4 weeks recovery, mice were bled to monitor reconstitution and started on HFD treatment for an additional 20 weeks. To distinguish chimerism, transplantation studies made use of congenic C57BL/6 strains carrying functionally equivalent alleles of the pan leukocyte marker CD45. For studies with PPARγ^+/+^ recipient mice, mice harboring the CD45.1 allele were purchased from Jackson and transplanted with bone marrow from PPARγ^+/+^ or PPARγ^A/A^ mice, which carry the wildtype allele for CD45 (CD45.2). For studies with PPARγ^A/A^ recipient mice, PPARγ^A/A^ mice were transplanted with bone marrow from PPARγ^+/+^ or PPARγ^A/A^ mice heterozygous for CD45.1.

### Statistical analysis

Data were analyzed using Prism software (GraphPad Software, Inc) and are expressed as mean ± SEM. Two-tailed Student’s *t* tests, one-way ANOVA with Newman-Keuls Multiple Comparison test, and two-way ANOVA were used to compare means between groups as indicated; *P* < 0.05 was considered significant.

## Supplementary Information

**Supplemental Figure 1.**
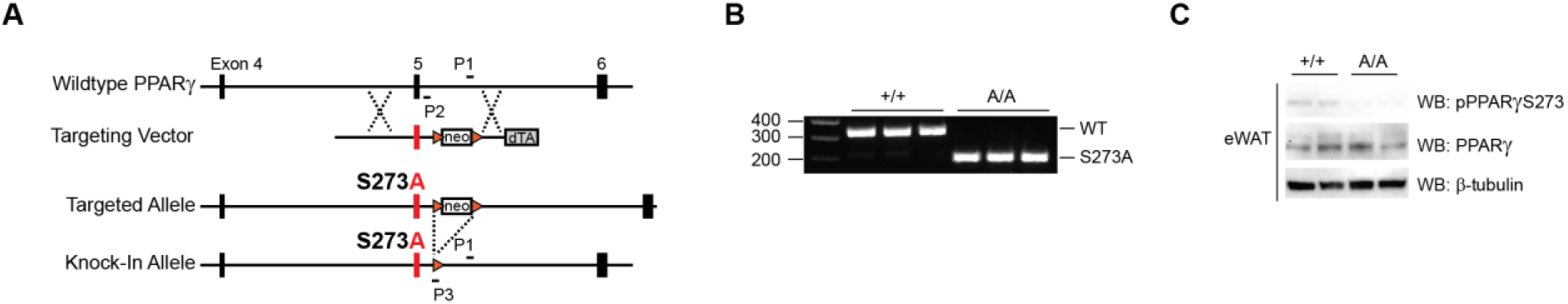
Generation of PPARγS273A knock-in mice. A. Targeting strategy for generation of PPARγ S273A knock-in mice representing the WT locus, targeting vector design, correctly targeted allele, and targeted allele following Cre-mediated recombination in the germline. Resulting knock-in mice contain an alanine substitution in both PPARγ isoforms (PPARγ2S273Aand PPARγlS243A). Orange triangles represent loxP sites. PCR primers for genotyping are indicated. B. Representative genotyping PCR results from tail DNA of wildtype and homozygous knockin mice performed with primers P1, P2, and P3. C. Immunoblot analysis reveals unphosphorylated PPARγS273 in epididymal adipose tissue (eWAT) of mice homozygous for PPARγS273A (A/A).

**Supplemental Figure 2.**
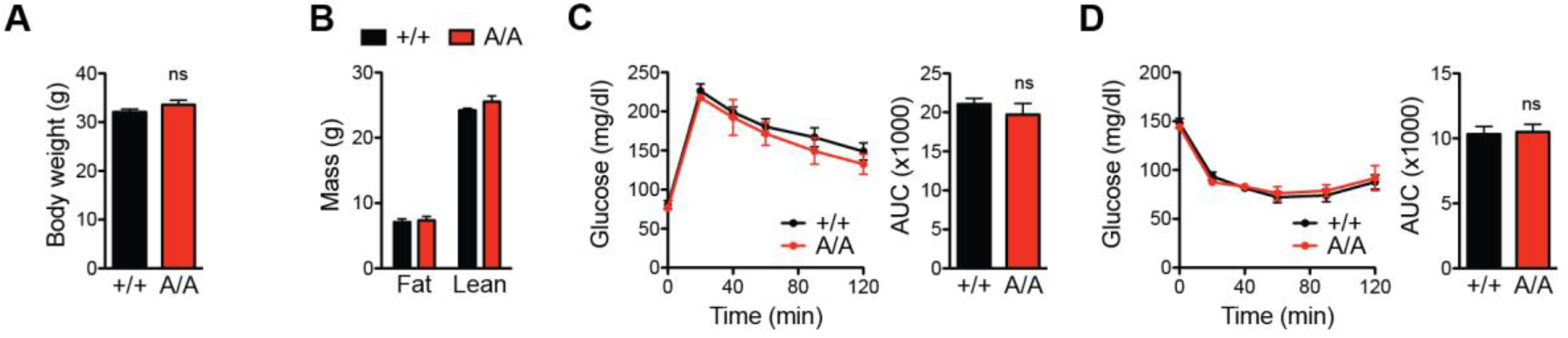
Phosphorylation is required for phenotypic differences in PPARγA/A mice. Metabolic profiling of wildtype (+/+) and PPARγA/A (A/A) mice maintained on a standard chow diet. A. Body weight of mice at 25 weeks of age (n = 7 +/+, 6 A/A), an age-match for mice in Figure 1. B. Body composition. C. Glucose tolerance test (1 g glucose/kg body weight; n = 4 +/+, 3 A/A). D. Insulin tolerance test (1.5 U insulin/kg body weight; n = 7 +/+, 5 A/A). C-D. Right, area under the curve (AUC) for the respective test. With the exception of C, data are pooled from two independent cohorts and presented as mean ± SEM; ns, not significant by Student’s t test.

**Supplemental Figure 3.**
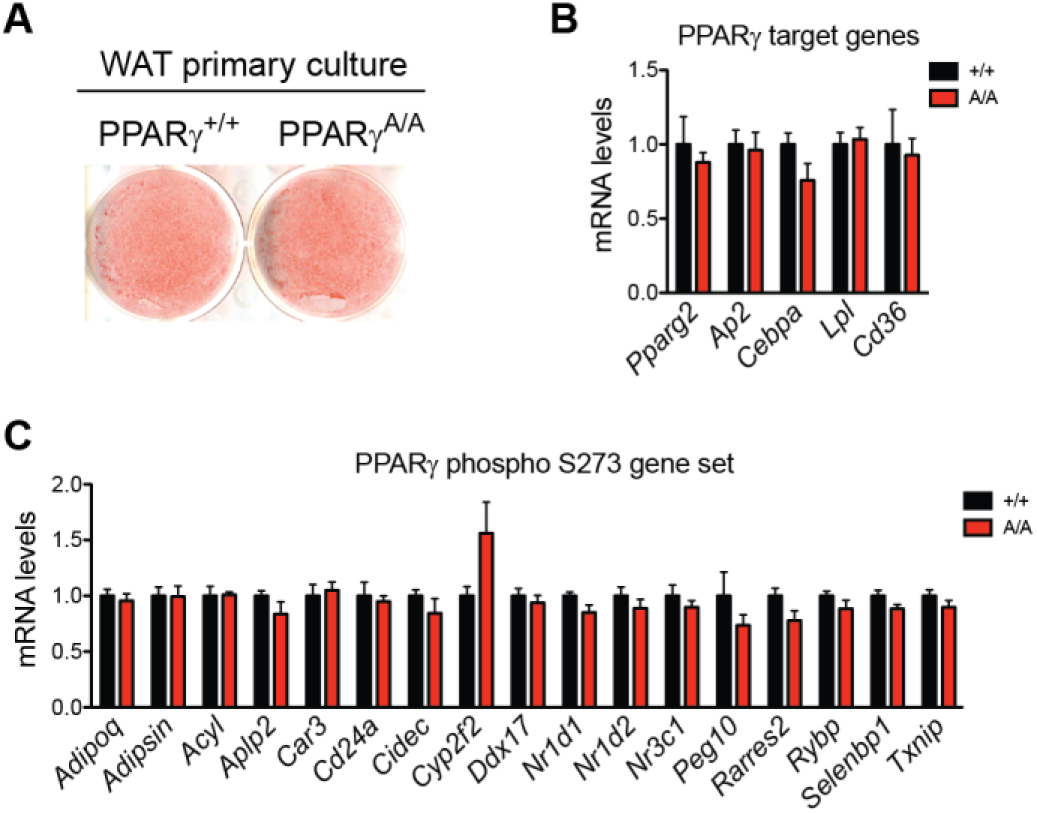
Cell-autonomous characterization of PPARγA/A adipocytes. A. Lipid accumulation of wildtype (+/+) and PPARγA/A (A/A) adipocytes differentiated ex vivo as shown by Oil Red O staining of representative images. B-C. mRNA levels in these adipocytes were determined for (B) PPARγ target genes and (C) genes previously shown to be sensitive to PPARγ phosphorylation at Ser273. Data are presented as mean ± SEM; no significant differences were observed by Student’s t test.

**Supplemental Figure 4.**
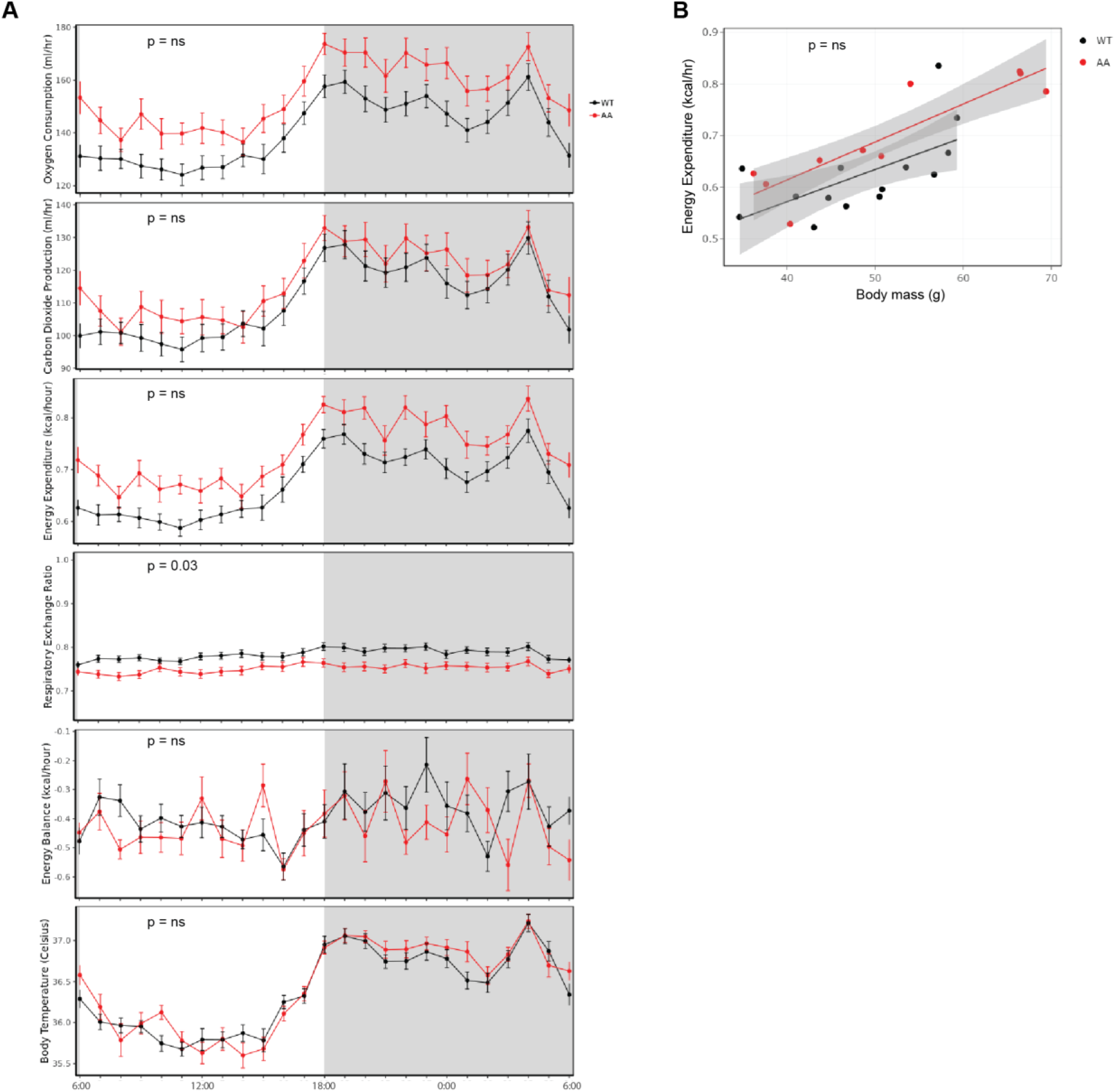
Energy Balance in PPARγA/A (A/A) mice. A. Indirect calorimetry measurements of wildtype (+/+, n = 6) and A/A (n = 5) mice maintained on a HFD. Shown are plots for oxygen consumption, carbon dioxide production, energy expenditure, respiratory exchange ratio, energy balance, and core body temperature after a 48-h acclimatization period. Shaded areas indicate the dark phase of the light cycle. Data are presented as mean ± SEM; ns, not significant; by ANOVA (RER and Energy Balance) or ANCOVA (others). B. Regression plots of energy expenditure vs total body mass.

**Supplemental Figure 5.**
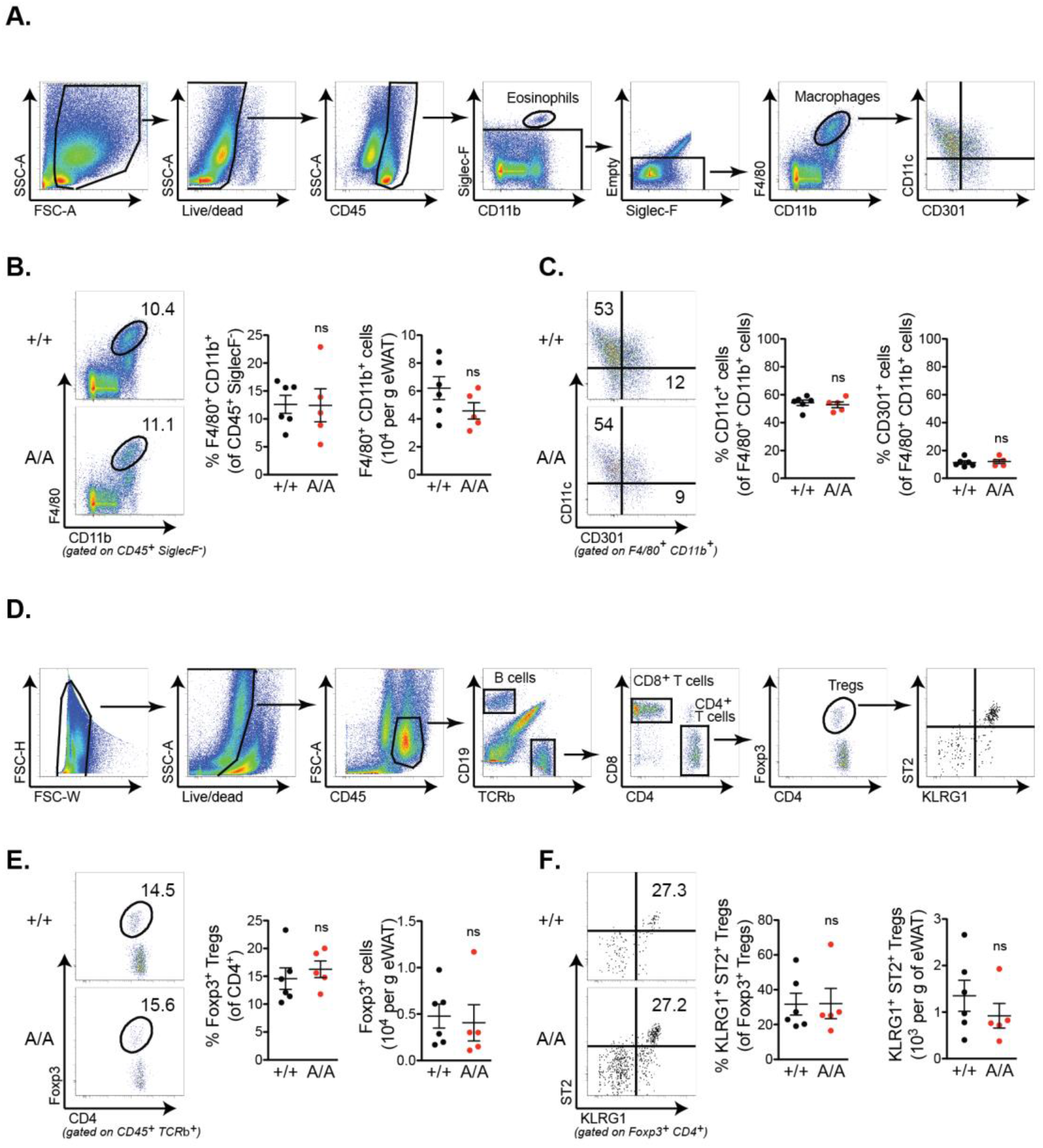
Immune cell populations are unchanged in adipose tissue of PPARγA/A mice after HFD feeding. A. Gating strategy for analysis of myeloid cells in epididymal adipose tissue (eWAT). Mononuclear cells from eWAT SVF were gated for forward and side-scatter (FSC/SSC) and live/dead prior to identification of hematopoietic cells (CD45^+^). After exclusion of eosinophils, CD45^+^ SiglecF-F4/80^+^ CD11b^+^ cells were defined as macrophages. Macrophage subsets were identified as being CD11C^+^ or CD301^+^. B. Quantification of macrophages in eWAT of mice fed a high fat diet (HFD) for 16 weeks (n = 6 +/+, 5 A/A). Left, representative dot plots; center, summary data (for fraction of CD45^+^ SiglecF^−^ cells); right, cell numbers per g of tissue. C. Macrophage subsets were identified as being CD11C^+^ or CD301^+^ and presented as fraction of F4/80^+^ CD11b^+^ cells. D. Gating strategy for analysis of lymphoid cells in eWAT SVF included exclusion of aggregates, live/dead gating, and identification of CD45^+^ CD19^+^ B cells, CD45^+^ TCR β^+^ CD8^+^ T cells, CD45^+^ TCR β^+^ CD4^+^ T cells, and CD45^+^ TCR β^+^ CD4^+^ Foxp3^+^ Treg cells. E. Quantification of Treg cells are presented as fraction of CD4^+^ T cells and per g of tissue. F. Treg cells expressing KLRG1 and ST2 are quantified as fraction of Foxp3^+^ Treg cells and per g of tissue. Data are presented as mean ± SEM; ns, not significant by Student’s t test.

**Table S1.**
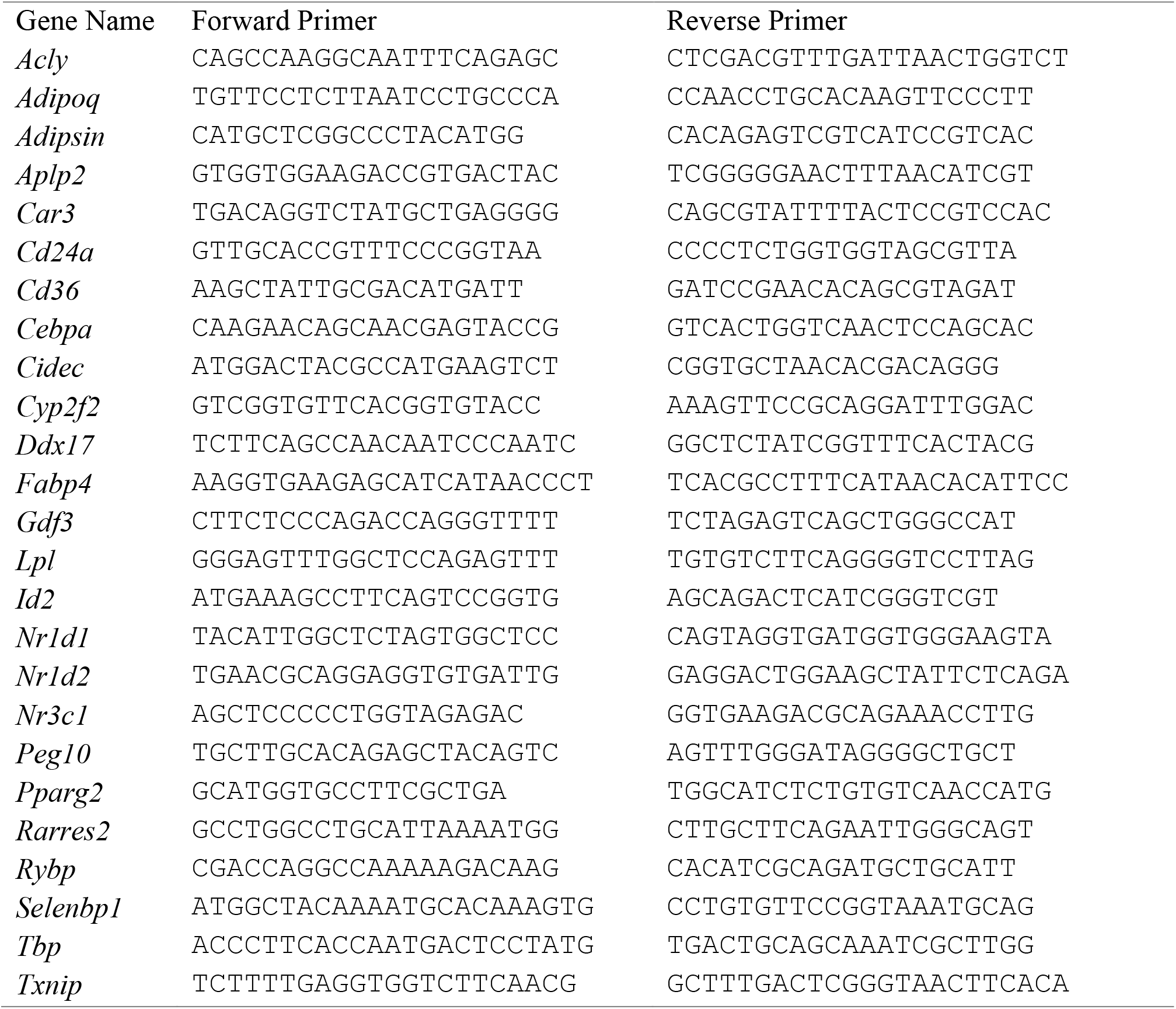
Primer sets used in this study

## Acknowledgements

Financial support for this project was provided from the NIH to ASB, R01DK107717, P30DK34854, and U24DK076169 for the NIDDK Mouse Metabolic Phenotyping Centers under the MICROMouse Program. JAH received support from the NIH T32DK007529-28. DR was supported by the Swiss National Science Foundation’s Early Postdoc Mobility grant. EDR was supported by NIH R01 DK102173 and DK113669. TB was supported by the São Paulo Research Foundation (Postdoc financial support #2017/25881-0). RNA-seq was performed by the Boston Nutrition and Obesity Research Center’s Functional Genomics and Bioinformatics Core (BNORC, NIH 5P30DK046200).

